# Neural and Gait Adaptations to Dual-Task Demands Across Development

**DOI:** 10.64898/2026.10.26.761688

**Authors:** Paige R. Nicklas, Edward G. Freedman, John J. Foxe

## Abstract

Cognitive-motor interactions describe the complex interplay underlying how the brain manages concurrent demands on motor and cognitive processing. Yet, how cognitive-motor interactions present across childhood and young adulthood remains understudied, particularly through multi-modal methods which include neurophysiology. We investigated development of cognitive-motor interactions using a dual-task study design including simultaneous motor (walking) and cognitive (visual Go-NoGo) tasks. Using mobile brain-body imaging, (MoBI), we recorded simultaneous EEG, gait kinematics, and task performance. 61 neurotypical 8-to-30-year-olds completed the Go-NoGo while seated or walking on a treadmill. With increased age, participants became faster and more accurate on the task, walking became less variable, and N200 and P300 ERP amplitudes increased. Two distinct effects of cognitive-motor interactions emerged with age. An N200 amplitude attenuation when walking became exaggerated with age, suggesting older participants engaged greater neurophysiological flexibility to support their improved task performances, whereas children performed less well and may not have needed to recruit the same adaptive neural mechanisms. Additionally, an attenuation of stride variability during dual-task walking lessened with age, demonstrating robust motor development under dual-task conditions. These findings demonstrate a distinct pattern of changes to cognitive-motor interactions through development. Future research can examine how this pattern varies in different developmental conditions.

## 1. Introduction

Almost all daily activities require combining cognition with movement for successful execution. Navigating grocery aisles with your cart while remembering items you need to buy is just one example of situations in which our brains are taxed across both cognitive and motor domains. This requires dynamic coordination of neural resources to successfully complete simultaneous tasks, but there are limitations to the complexity or number of concurrent tasks that the brain is capable of coordinating. Understanding the developmental trajectory of how individuals acquire the capacity to effectively coordinate simultaneous cognitive and motor demands is critical for understanding how these skills emerge, mature, and ultimately shape everyday functioning.

Commonly referred to as cognitive-motor (C-M) interference (Al-Yahya et al., 2011), performance declines are often observed when people engage in multiple tasks at the same time (“dual-tasking” or “multi-tasking”)(Pashler, 1994). However, and perhaps counter-intuitively, studies have also shown mixed effects, wherein C-M dual-task conditions can result in maintenance or even improvement in behavioral task performance (De Sanctis et al., 2014; De Sanctis et al., 2020; Lim et al., 2025; Malcolm et al., 2015; Malcolm et al., 2019; Patelaki, Foxe, Mantel, et al., 2023; Patelaki et al., 2022; Patelaki, Foxe, McFerren, et al., 2023). Declines in task performance or motor parameters observed when comparing dual-tasking to single-tasking are often referred to in the literature as “dual-task costs”, although this terminology reflects the historical emphasis on performance decrements despite the growing literature that dual-task “benefits” can also occur. Current evidence supports framing C-M interactions as a spectrum and using the more encompassing term: “interactions”, since individuals can exist anywhere between strong interference and strong integrative effects. A common way to study these C-M interactions is with dual-task experimental designs, wherein performance on a cognitive task alone (single-task) is compared to performance on the same task when a simultaneous motor demand is introduced, such as walking. The motor task is also assessed alone to form a baseline for parameters of interest and then compared against the dual-task condition (Huang & Mercer, 2001; Leone et al., 2017).

To gain insight into the neurophysiological activity underlying C-M interactions, EEG-based Mobile Brain/Body Imaging (MoBI) methods allow assessment and comparison of brain activity between single and dual-task conditions. Research in this area is broad in design such as exact choice of tasks, dependent measures, and techniques, but all generally support the concept of capacity-limited neurological resources and ability to reallocate them as needed influencing the extent of experienced dual-task costs (or benefits), and the idea that the flexibility to deploy these resources declines from young to older adulthood (Belur, 2020; De Sanctis et al., 2020; Krasovsky, 2018; Patelaki et al., 2022; Richardson et al., 2022; Ward et al., 2022). Collectively, the literature provides evidence of variations in the impacts of dual-task paradigms and shows that some individuals may have more neural resources at their disposal, and more effective neural flexibility to distribute those resources. Notably though, these studies have been thus far limited to adults and aging populations. A major remaining question is how the ability to adapt to multi-modal demands presents during childhood and develops through adolescence and young adulthood, how the underlying neurophysiology presents, and by extension, if characteristics of C-M interaction can serve as informative markers for research, clinical, and educational applications across development.

Some dual-task studies in children (Cinar et al., 2021; Hagmann-von Arx et al.,2016; Hocking et al., 2020; Möhring et al., 2021; Rabaglietti et al., 2019; Saxena et al., 2019; Schott & Klotzbier, 2018; Subara-Zukic et al., 2024), generally support the notion that, as with most functions (Hartshorne & Germine, 2015), the ability to successfully and efficiently coordinate demands across functional domains improves with age as the brain and body mature. However, a vital gap in these studies is the lack of neuroimaging and of methods capable of capturing real-time changes synchronized across domains. There is a need to understand the neuroscience behind the development of C-M interactions. While dual-tasking and its neural underpinnings have been well-studied in adults, (for C-M neuroimaging reviews, see Richer et al., 2024(Richer et al., 2024); Leone et al., 2017(Leone et al., 2017), Gramann et al., 2014(Gramann et al., 2014)), to our knowledge, there is only one other mobile EEG study conducted in children using a C-M dual-task design (Fauvel, 2025). That group used an auditory oddball task and treadmill walking in children aged 7-12 and in adults. They found that “attentional resource management” improved with age, with older children showing increased task performance. When comparing motion states, sitting and walking, they found that cognitive performance did not change, but neurophysiology was attenuated when walking in the children, whereas these changes were not observed in their adults. Taken together, their results support the notion that C-M interactions and the ability to efficiently manage them improves through childhood and adolescence, and emphasize the need to deepen our understanding of the dynamic and complex interplay of these interactions across development. To our knowledge, only one study specifically using MoBI has included participants under 18 years of age (Nicklas et al., 2026), although this was an analysis of a group of adolescents aged 13-23 years, and individual age was not incorporated into the analyses.

Here, we set out to use age as a continuous variable in a neurotypical cohort to establish better understanding of C-M interactions and neurophysiology across childhood and into early adulthood. We used MoBI with a visual Go-NoGo (vGNG) response inhibition task. The vGNG was chosen for a few reasons. Young children are capable of understanding and completing response inhibition tasks such as this one (Barry et al., 2022; Jonkman, 2006), as executive functions required to do so develop early in childhood (Best & Miller, 2010). More broadly, the vGNG task is also well-established and its associated neurophysiology is robustly understood, including the N2 and P3 event-related potentials (ERPs) (Falkenstein et al., 1999; Huster et al., 2013; Luck, 2014). The N200 is a negative voltage deflection understood to reflect cognitive attention and control, stimulus distinction, and conflict monitoring. It is fronto-centrally distributed on the scalp, peaks about 200-350 ms after the stimulus (Folstein & Van Petten, 2008), and is associated with generators in the anterior cingulate cortex (Bekker et al., 2005; Brydges et al., 2013; Dias et al., 2003; Dias et al., 2006; Iannaccone et al., 2015). The P300 reflects the decision-making needed to complete the assigned task (Falkenstein et al., 1999; Malcolm et al., 2015; Malcolm et al., 2019; Patelaki et al., 2022), arrives approximately 150 ms after the N200, is a positive voltage deflection observed more centro-parietally and peaks 350-600 ms after stimulus onset. It is understood to have a wider distribution of underlying neural generators, likely because it’s thought to reflect parallel processing occurring within both cognitive and motor circuits for complex processes such as stimulus evaluation, motor planning, and memory updating.(Huster et al., 2013; Kirmizi-Alsan et al., 2006; Patelaki et al., 2022). Lastly, this specific task also has well-established characteristics specifically within the MoBI literature (De Sanctis et al., 2012; De Sanctis et al., 2014; De Sanctis et al., 2020; Malcolm et al., 2015; Malcolm et al., 2019; Patelaki, Foxe, Mantel, et al., 2023; Patelaki et al., 2022; Patelaki, Foxe, McFerren, et al., 2023).

Our central hypothesis is that C-M interactions are more interfering at younger ages, which will be reflected in the magnitude of dual-task costs. This greater interference should be evidenced through MoBI by weaker ERP differences between sitting and walking, worsening of cognitive task performance when walking compared to sitting, and weaker adjustments of gait kinematics when walking by itself compared to walking while doing the cognitive task.

## 2. Methods & Materials

### 2.1 Participants

35 young adults (18-30 years old, mean age = 22.61 ± 3.15 years, 15 male, 20 female) and 26 children (8-17 years old, mean age = 12.65 ± 2.82 years, 8 male, 18 female) participated in this study. All participants provided written informed consent or, if under 18 years old, provided informed assent and their parents/guardians provided consent. All participants reported no diagnosed neurological or neurodevelopmental conditions, no recent head injuries, and normal or corrected-to-normal vision. All study procedures were approved by the University of Rochester Institutional Review Board (STUDY00006574). Participants were paid $18 per hour for time spent in the lab. Parents/guardians accompanying participants under the age of 18 were also given a flat $20 payment for each visit. The age groups were matched on full-scale intelligence quotient (FSIQ), verbal-IQ, and nonverbal-IQ. Two children and one adult did not attend the study session for assessment, so they are excluded from IQ data. One adult had experience administering the IQ test used, the Wechsler Abbreviated Scale of Intelligence, 2nd Edition (WASI-II), and therefore was not given the assessment. This creates group sizes of 33 young adults and 24 children for IQ. Descriptives and independent samples t-tests are shown in Table 1.

**Table 1.**
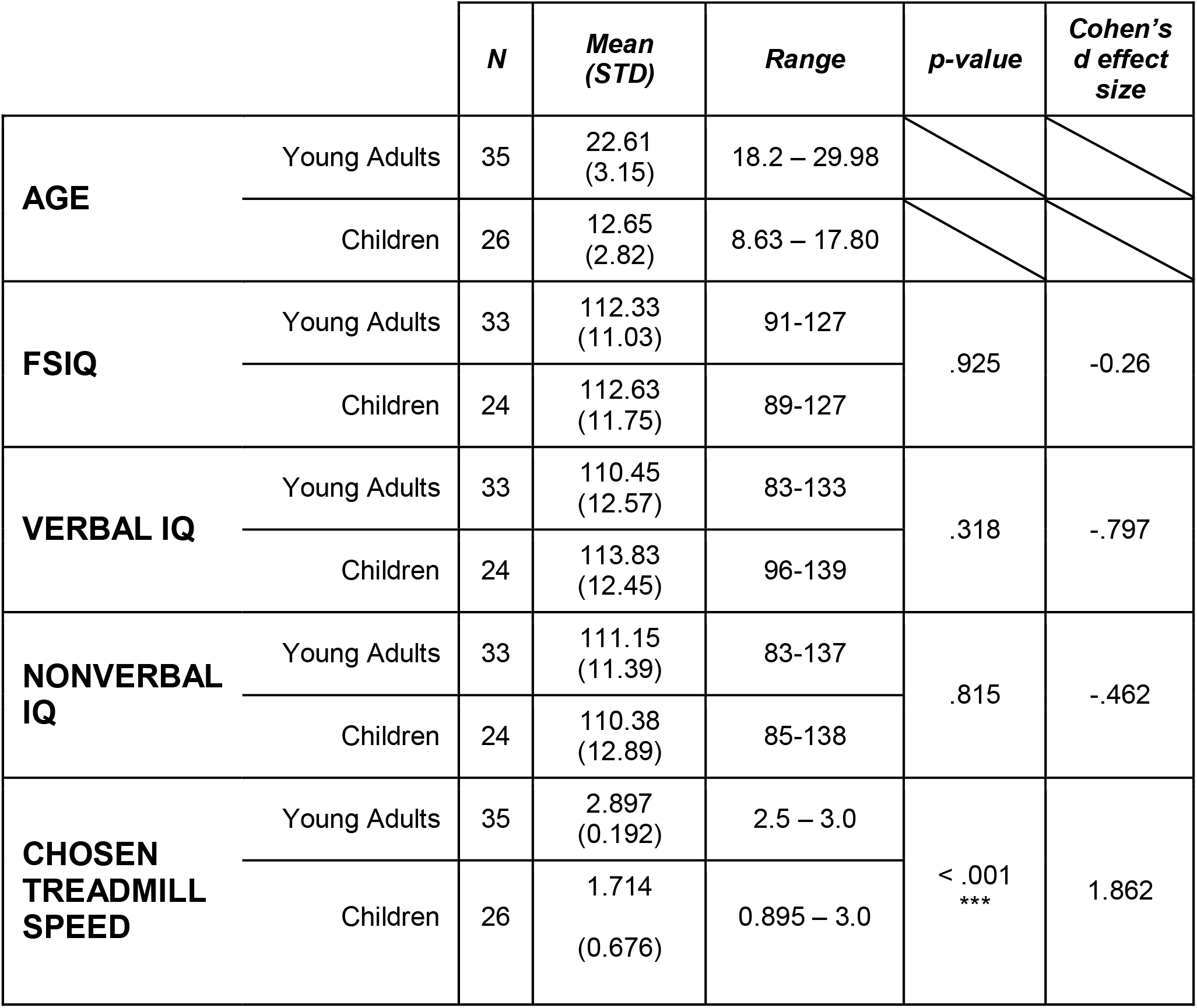
Group descriptives and independent samples t-tests.

### 2.2 Experimental Procedure and Cognitive Task

The study involved participants performing a cognitive task while either seated, or walking on a treadmill (Tuff Tread, Conroe, TX, USA). The experiment consisted of 16 total blocks, of which the 1^st^ was an abbreviated (approximately 1 minute) seated training block, followed by 7 sitting, 7 walking, and 1 walking-only block(s) presented in a pseudorandom order, such that no more than 2 blocks of the same motion state could occur in succession. Each block was 4-5 minutes long and participants took a break between each block. During breaks participants were provided with drinks/snacks whenever requested. The training block was completed first to ensure each participant understood the task instructions and could be repeated if necessary. Response accuracy feedback was not given to participants. Treadmill walking speed was determined collaboratively between the participant and research team. Participants were instructed that the walking speed should feel like a brisk walk, but they should not feel like they needed to transition to a jog. Further, participants were instructed to walk as normally as possible, to only hold onto side safety rails when needed, and to let their arms swing naturally.

Participants completed a visual Go-NoGo (vGNG) response inhibition task during all experiment blocks, except for the walking-only block. Each block consisted of 240 trials (approximately 4 minutes), wherein 87% (209) were “Go” trials and 13% (31) were “NoGo” trials. Within a single trial, stimuli were presented for 67 ms, followed by an image of a white fixation cross for the remainder of the trial. Stimulus offset asynchrony was 1017 ms. Responses given in the first 183 ms post-stimulus-onset were considered delayed responses from the previous trial and were not included as a correct response. These parameters are as presented originally in Patelaki et al., 2022 (Patelaki et al., 2022). One key difference between the original task and the task used in the current study is the stimulus set included. The stimulus pictures were taken from the International Affective Picture System (IAPS) database (Bradley & Lang, 2017). Only positive- and neutral-valence pictures were used in the current study (i.e. negatively valenced images were not employed).

Stimulus presentation was programmed using Presentation software version 21.1 (Neurobehavioral Systems, Albany, CA, USA) and either projected (InFocus XS1 DLP, 1024 x 768 pxl) centrally onto a projection screen approximately 1.2 meters in front of the treadmill, or on a TV screen (LG - 83” Class C3 Series OLED evo 4K UHD Smart webOS TV, Model: OLED83C3PUA). During the walking-only block, the white fixation cross was on the screen for the full block duration. Participants held a Nintendo Switch™ Joy-Con controller (Nintendo of America Inc., Redmond, Washington, USA) in their dominant hand to give responses to the task. All participants were instructed to compare every picture to the one that immediately preceded it. If the picture was different, they were to press the button as quickly as they could (Go Trial). However, if the photo shown was exactly the same as the one just before it, participants were instructed to withhold their button press (NoGo Trial), (Figure 1). Study personnel confirmed that each participant continued to understand the task instructions throughout the blocks via verbal confirmation (e.g., asking “What should you do if you see 2 photos in a row that are exactly the same?”), and real-time monitoring of responses. Time-synchronized stimulus triggers for the vGNG task, button responses, EEG recording, and motion-tracking were collected using Lab Streaming Layer (LSL) software (Swartz Center for Computational Neuroscience, University of California, San Diego, CA; https://github.com/sccn/labstreaminglayer).

**Figure 1.**
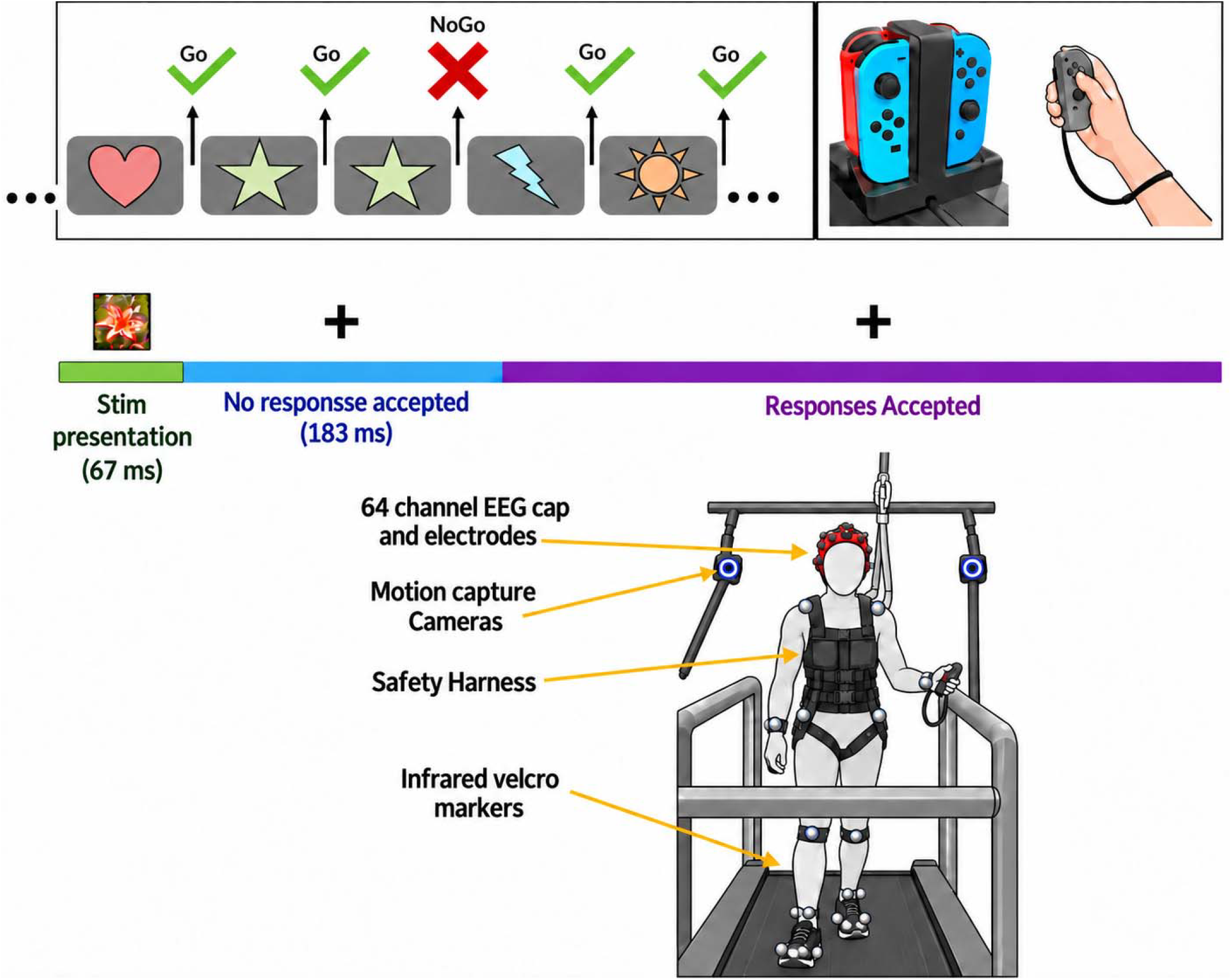
Schematic of visual Go-NoGo cognitive task and participant set-up

**Figure 2.**
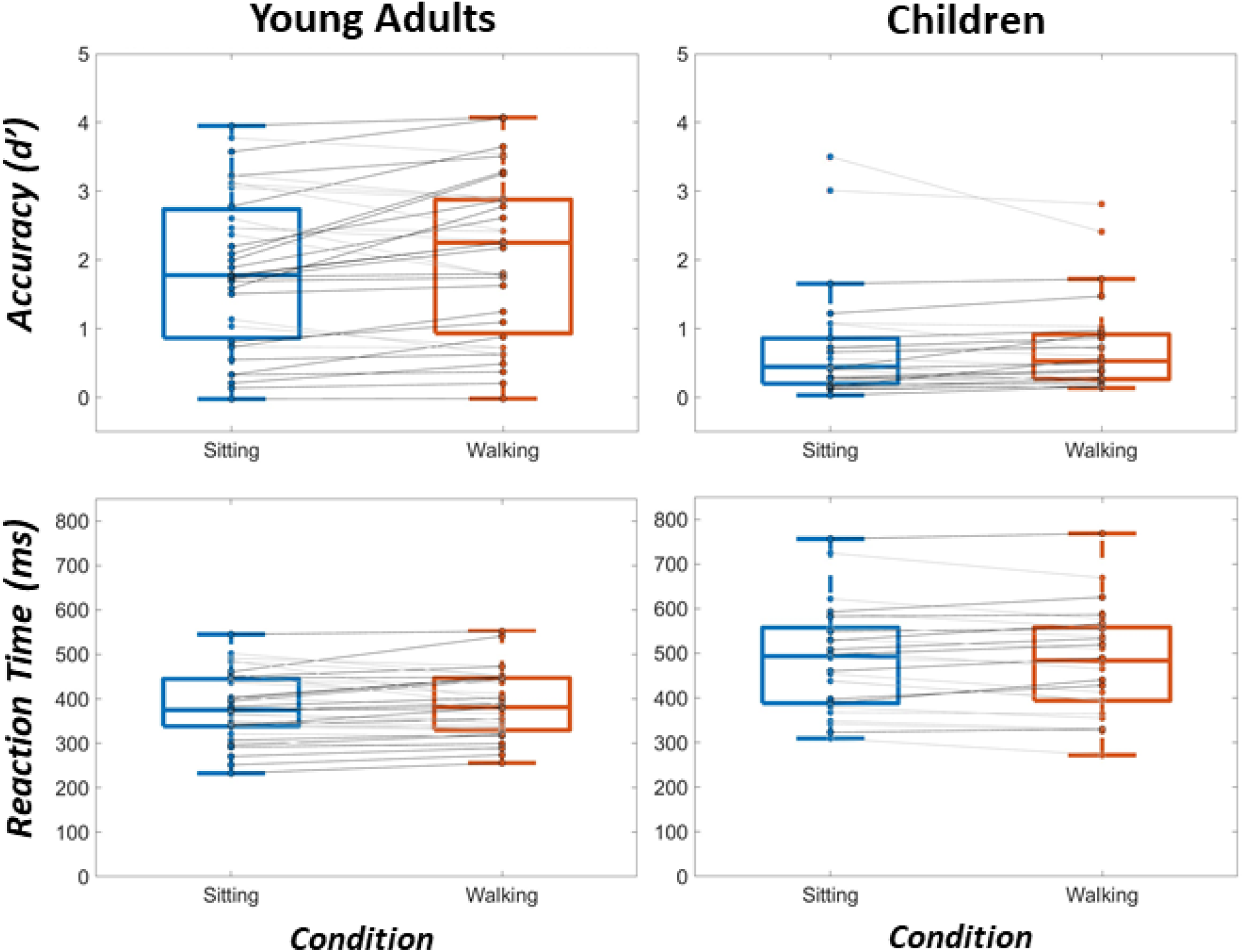
Response accuracy and reaction time on the visual Go-NoGo task. Left column is Young Adults, right column is Children. Top row is Accuracy (d’) and bottom row is Reaction Time (ms). Within each plot, the blue boxplot corresponds to the sitting condition, and red corresponds to walking. Each dot on the boxplots is a single participant. A single participant’s sitting value is connected to their respective walking value by a gray line. The center line of each boxplot is the median, the top and bottom edges of the boxes are the 75th and 25th percentiles, respectively, and the top and bottom whiskers are the most extreme data points not considered outliers.

### 2.3 Task Performance Measures & Analyses

The cognitive task instructions create four possible behavioral outcomes: 1) Hits, when a button response is correctly given during a Go trial; 2) Misses: when a button response is incorrectly withheld during a Go trial; 3) False Alarms: when a button response is incorrectly given during a NoGo trial; and 4) Correct Rejections: when a button press is correctly withheld during NoGo trials. The dependent measures recorded from these outcomes are reaction time during Hits, and task accuracy. Task accuracy is reported as d’, a sensitivity index [d’ = *z*(False Alarm Rate) – *z*(Hit Rate)](Green & Swets, 1966). Statistical analyses for task performance measures were conducted using repeated measures ANCOVAs with the within-subjects factor of motion state (sitting versus walking), and age as a continuous covariate. If a main effect of the within-subjects factor of motion state is found, Bonferroni’s correction was used for *post-hoc* analyses. Significance threshold was set at *α* = 0.05. Main effects or interaction effects involving the continuous covariate of age cannot be further probed with typical *post-hoc* testing, so these effects are further probed with linear regressions but interpreted carefully in the discussion due to lack of ability to control for multiple comparisons.

### 2.4 EEG Recording, Processing, and Visualization

Electroencephalography (EEG) was recorded from 64 channels with the BioSemi ActiveTwo amplifiers (BioSemi, Amsterdam, the Netherlands) with an anti-aliasing filter (−3 dB at 3.6 kHz) following the international 10–20 system. The system uses active electrodes and a direct current (DC) coupling with a hardware bandpass filter from DC to 150 Hz to reduce low-frequency drift and high-frequency noise. The recording reference was the Common Mode Sense (CMS) active electrode, which, together with the Driven Right Leg (DRL) passive electrode, forms a feedback loop to stabilize the reference signal and reduce common-mode noise. This referencing convention in BioSemi systems differs from traditional reference electrodes by dynamically compensating for potential differences between the scalp and the amplifier ground, effectively creating a “zero potential” point. The sampling frequency was 512 Hz.

All EEG processing and analyses were performed using custom MATLAB scripts (MathWorks Inc., Natick, MA, United States) and functions from EEGLAB (Delorme & Makeig, 2004) or FieldTrip (Oostenveld et al., 2011) toolboxes. Continuous EEG data were parallel filtered at a bandpass of 0.1-45 Hz and of 1-45 Hz, applied using zero-phase filtering (“filtfilt” in EEGLAB) and hamming window types. The average reference was used. Bad channels were automatically detected and manually confirmed, then interpolated using the spherical method. Trials were sorted based on type, Go or NoGo, and motion state, sitting or walking. For the N200 and P300 ERP components, epochs were time-locked to the onset of stimulus presentation and extended from 200 ms preceding stimulus onset (-200 ms) to 800 ms after, for a total epoch window of 1000 ms. Independent component analysis (ICA) was performed using the AMICA algorithm (adaptive mixture ICA) (Palmer, 2008; Sejnowski, 1996). AMICA decomposition is biased toward high-power, low-frequency data, and therefore for an optimal approach, the decomposition was performed on the dataset preprocessed from 1-45 Hz, and then the resultant spheres and weights are back-projected on to the dataset with the 0.1-45 Hz bandpass (Pion-Tonachini, 2019) that computes IC class probabilities for components. ICs with a brain classification higher than the summed probabilities of all other artifacts and where ‘Brain’ and ‘Other’ summed were greater than 0.5 were retained. Resultant ICs were inspected and classified using an automated component classifier (ICLabel v1.2.5).

Grand average ERPs are plotted by motion state (sitting or walking) and age group (Young Adults (YAs) and Children (Cs)). The waveforms plotted are from correct rejection (CR) trials, as these represent successful completion of the more cognitively demanding trial type and would most robustly capture C-M interactions. Grand average scalp topography plots are averaged in 10 ms increments across the 50 ms centered on that age group’s overall mean peak latency for each motion state. For the N200, mean peak latencies at FCz were 261 ms for YAs and 239 ms for Cs. For the P300, mean peak latencies at CPz were 446 ms for YAs and 424 ms for Cs.

### 2.5 ERP Peak Analyses

Peak amplitudes and latencies were calculated for every subject for the N200 and P300 ERPs based on time-windows and electrode locations well-established in previous literature. For the N200, peaks were calculated based on the most negative value in the 200-325 ms time-window at FCz for each participant’s mean ERP (Bekker et al., 2005; Brydges et al., 2013; Falkenstein et al., 1999; Folstein & Van Petten, 2008; Iannaccone et al., 2015). For the P300, the most positive point within the time-window of 400-650 ms at CPz was calculated (Falkenstein et al., 1999; Huster et al., 2013; Kirmizi-Alsan et al., 2006; Malcolm et al., 2015; Malcolm et al., 2019; Patelaki et al., 2022). We chose to plot and calculate peaks at specific electrode locations instead of averaging across spatial clusters (i.e. “frontal-central cluster” or similar). The loci of the N200 and P300 ERPs are robustly characterized and spatially close. We did not want to create clusters within which these ERPs would overlap and be averaged together. Study personnel confirmed the chosen latency windows and channels visually using the group grand averages. A repeated-measures ANCOVA with a within-subject factor of motion state (sitting or walking) and a continuous covariate of age was performed using IBM SPSS Statistics version 29. If a main effect of motion state (walking or sitting) was found, *post-hoc* pairwise comparisons were conducted using Bonferroni’s correction. If a main effect of the covariate was found and/or an interaction between the covariate and motion state, linear regressions were used to further characterize the effect.

### 2.6 Exploratory Statistical Cluster Analyses of EEG Data

To further explore this rich high-density dataset, *post-hoc* statistical t-value cluster-based permutation analyses were also used. Pointwise two-tailed paired t-tests were applied across all samples of the epoch window for all electrodes. A cluster was considered significant only if consecutive points exceeded the set α = 0.05 threshold for a minimum of 10 consecutive sample points (De Sanctis et al., 2014; Foxe & Simpson, 2002; Guthrie & Buchwald, 1991; Patelaki et al., 2022; Richardson et al., 2022). As an exploratory approach, the cluster analyses are complementary to the a priori statistical analyses that only consider data from specified latency windows and channel locations. They are specifically deployed in a hypothesis-generating context, intended to generate hypotheses for future testing.

### 2.7 Gait Kinematic Recording & Analyses

The trajectories of the body-markers were recorded by the OptiTrack system, an optical 3D motion analysis system including 16 cameras that electronically connect to Motive software that collect data at 360 Hz (Optitrack Prime 41 cameras, Motive v2.1 software, Natural Point Corvallis, OR, USA). A total of 41 reflective markers were placed over anatomical landmarks (“Baseline + Hinged Toe” default skeleton marker set) on full-body, motion-capture suits worn by participants (Figure 1). For dependent variables of gait, the 4 markers placed on each foot (posterior of the calcaneus, tip of the second distal phalanx, distal end of the first metatarsal, and distal end of the fifth metatarsal) were used to measure stride distance, stride distance variability, stride time, and stride time variability. Participants also wore a custom-designed safety harness at all times, which was anchored into the ceiling. One child and three young adults do not have kinematic data due to technical issues with the motion-tracking software or poor recording quality, and are therefore excluded from these analyses. This makes the group sizes for these analyses 32 young adults, and 25 children. To process the gait data and calculate dependent measures, we first defined gait cycles for each participant using the heel markers. The axes of the 3 movement dimensions were defined as follows: X is the dimension of lateral movement (right/left relative to the motion of the treadmill belt), Y represents anterior-posterior movement (parallel to the motion of the treadmill belt), and Z is the dimension of vertical lift movement (up-and-down relative to the motion of the treadmill belt). We used the time series of X, Y, Z coordinates for each foot to determine heel strikes. A heel strike was defined as the local maxima of the Y position timeseries. Then, a gait cycle was defined as the period between 2 consecutive heel strikes of the same foot. Further, our processing pipeline excluded gait cycles with a kurtosis exceeding 5 standard deviations of the mean. Strides were then calculated and defined as 2 consecutive gait cycles of alternating feet (one step of the right foot followed by one step of the left foot). We calculated each participant’s mean stride distance and time, and the variability of each of those measures. For variability we used Coefficient of Variation (CV% = (SD/mean) × 100). A repeated-measures ANCOVA with a within-subject factor of dual-task state (walking only or walking while performing the cognitive task) and a continuous covariate of height was performed in SPSS. If a main effect of dual-task state was found, *post-hoc* pairwise comparisons were conducted using Bonferroni’s correction. If a main effect of the covariate was found and/or an interaction between the covariate and dual-task state, linear regressions were used to further investigate the effect.

## 3 Results

### 3.1 Participant Characteristics

### 3.2 Task Accuracy and Reaction Time

An rmANCOVA including a categorical within-subjects factor of motion state (sitting and walking) and a continuous covariate of age showed a significant main effect of age (*F*(1,59) = 29.325, *p* < .001, η_p_^2^ = 0.332), indicating that d’ increased with age. There was no significant main effect of motion state (*p* = .938), nor a significant interaction between age and motion state (*p* = .540). Similarly, reaction time (RT) showed only a main effect of age (*F*(1,59) = 20.156, *p* < .001, η_p_^2^ = 0.225), and no effect of motion state (*p* = .074) nor interaction between age and motion state (*p* = .090). RT speeded up with age. Table 2 summarizes all results.

**Table 2.**
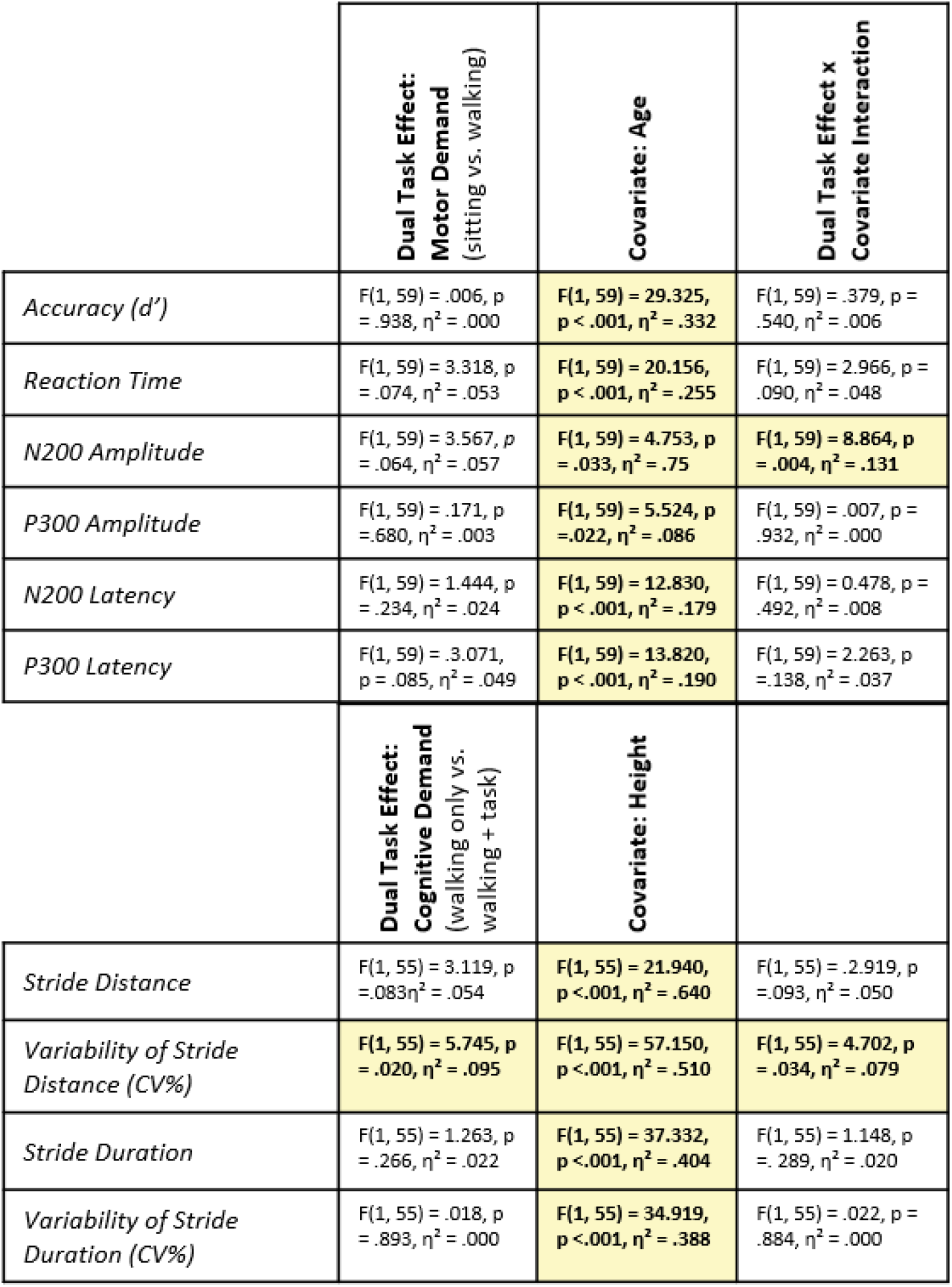
Summary of Results. Significant effects are filled with yellow and text is bolded.

|  | **Dual Task Effect:**<br>**Motor Demand**<br>(sitting vs. walking) | **Covariate: Age** | **Dual Task Effect x**<br>**Covariate Interaction** |
| --- | --- | --- | --- |
| *Accuracy (d')* | $F(1, 59) = .006, p = .938, \eta^2 = .000$ | **$F(1, 59) = 29.325, p < .001, \eta^2 = .332$** | $F(1, 59) = .379, p = .540, \eta^2 = .006$ |
| *Reaction Time* | $F(1, 59) = 3.318, p = .074, \eta^2 = .053$ | **$F(1, 59) = 20.156, p < .001, \eta^2 = .255$** | $F(1, 59) = 2.966, p = .090, \eta^2 = .048$ |
| *N200 Amplitude* | $F(1, 59) = 3.567, p = .064, \eta^2 = .057$ | **$F(1, 59) = 4.753, p = .033, \eta^2 = .075$** | **$F(1, 59) = 8.864, p = .004, \eta^2 = .131$** |
| *P300 Amplitude* | $F(1, 59) = .171, p = .680, \eta^2 = .003$ | **$F(1, 59) = 5.524, p = .022, \eta^2 = .086$** | $F(1, 59) = .007, p = .932, \eta^2 = .000$ |
| *N200 Latency* | $F(1, 59) = 1.444, p = .234, \eta^2 = .024$ | **$F(1, 59) = 12.830, p < .001, \eta^2 = .179$** | $F(1, 59) = 0.478, p = .492, \eta^2 = .008$ |
| *P300 Latency* | $F(1, 59) = .3071, p = .085, \eta^2 = .049$ | **$F(1, 59) = 13.820, p < .001, \eta^2 = .190$** | $F(1, 59) = 2.263, p = .138, \eta^2 = .037$ |
|  | **Dual Task Effect:**<br>**Cognitive Demand**<br>(walking only vs. walking + task) | **Covariate: Height** |  |
| *Stride Distance* | $F(1, 55) = 3.119, p = .083, \eta^2 = .054$ | **$F(1, 55) = 21.940, p < .001, \eta^2 = .640$** | $F(1, 55) = .2919, p = .093, \eta^2 = .050$ |
| *Variability of Stride Distance (CV%)* | **$F(1, 55) = 5.745, p = .020, \eta^2 = .095$** | **$F(1, 55) = 57.150, p < .001, \eta^2 = .510$** | **$F(1, 55) = 4.702, p = .034, \eta^2 = .079$** |
| *Stride Duration* | $F(1, 55) = 1.263, p = .266, \eta^2 = .022$ | **$F(1, 55) = 37.332, p < .001, \eta^2 = .404$** | $F(1, 55) = 1.148, p = .289, \eta^2 = .020$ |
| *Variability of Stride Duration (CV%)* | $F(1, 55) = .018, p = .893, \eta^2 = .000$ | **$F(1, 55) = 34.919, p < .001, \eta^2 = .388$** | $F(1, 55) = .022, p = .884, \eta^2 = .000$ |

Since the only main effects found here are of the continuous covariate, age, no further *post-hoc* pairwise comparisons were conducted. However, to further characterize this developmental main effect, a linear regression was performed for each dependent variable with age (d’ sitting (*F*(1,59) = 26.663, *p* < .001, R^2^ = 0.311), d’ walking (*F*(1,59) = 28.955, *p* < .001, R^2^ = 0.329), RT sitting (*F*(1,59) = 22.602, *p* < .001, R^2^ = 0.277), and RT walking (*F*(1,59) = 16.816, *p* < .001, R^2^ = 0.221)) (Figure 3). These results indicate that, regardless of motion state, age significantly predicts task accuracy and reaction time, with d’ increasing and RT speeding up with increased age. We will interpret and discuss these results conservatively since this approach does not control for multiple comparisons.

**Figure 3.**
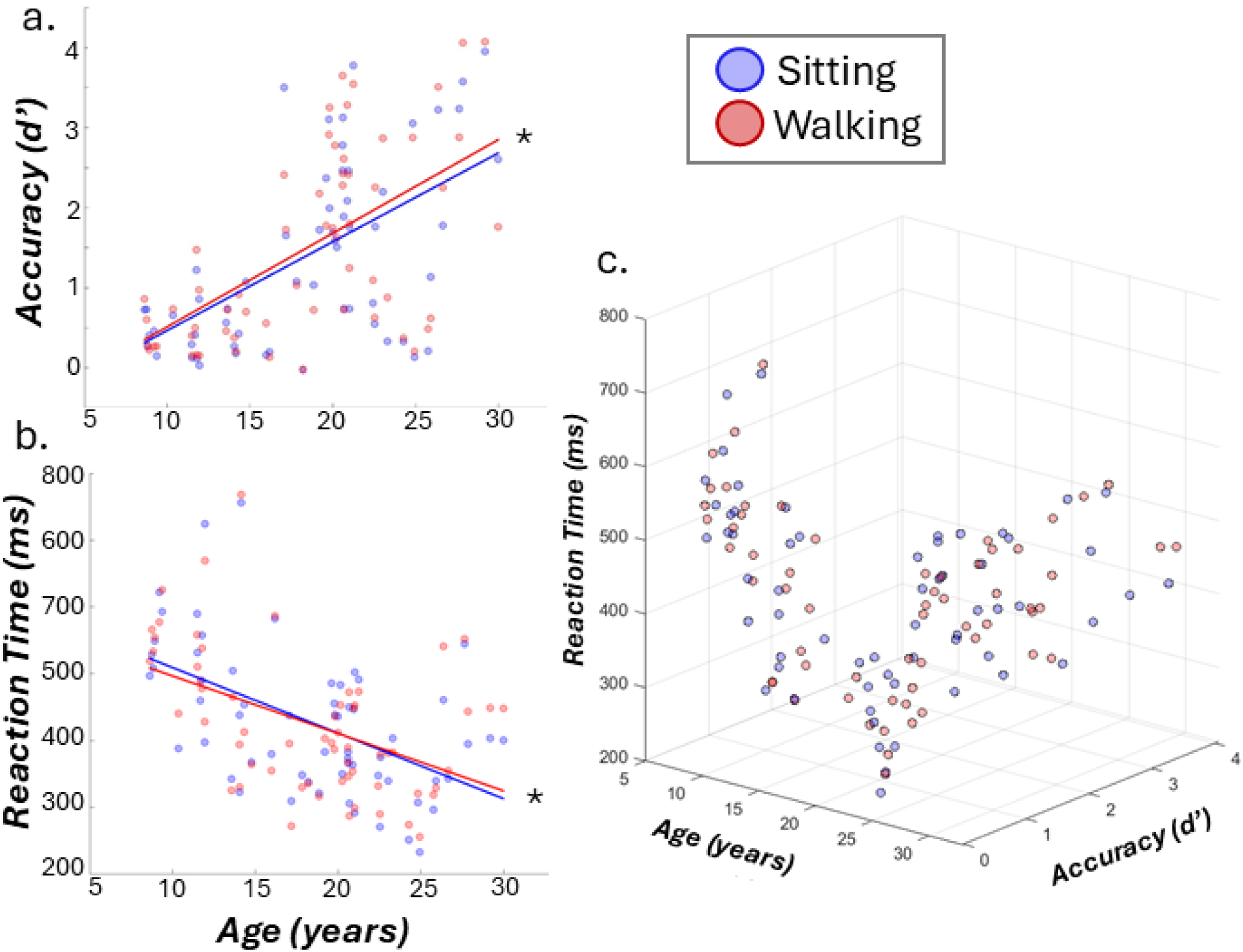
Characterization of the relationship between age and visual Go-NoGo task performance. A) Age and Task Accuracy (d’), B) Age and Reaction time (ms), and C) a 3D scatter plot of Age, Task Accuracy and Reaction Time. For all scatter plots, each dot is an individual participant and for each participant there is a blue dot for sitting and a red dot for walking. In A) and B) linear regression lines are plotted in blue for sitting and red for walking.

### 3.3 Neurophysiology

For the rmANCOVA on peak amplitudes and latencies, significant values are reported here along with partial-eta squared. If motion state has a significant main effect, Bonferroni’s *post-hoc* pair-wise tests were performed. However, there were no significant main effects of motion state. Results are visualized in Figure 4. All effects are summarized in Table 2. There was a significant effect of age for all peak measures: N200 peak amplitude (*F*(1,59) = 4.753, *p* = .033, η _p_^2^ = 0.075) and latency (*F*(1,59) = 12.830, *p* < .001, η_p_ ^2^ = .179); P300 peak amplitude (*F*(1,59) = 5.524, *p* = .022, η _p_^2^ = 0.086) and latency (*F*(1,59) = 13.820, *p* < .001, η_p_ ^2^ = 0.190). As shown in Figure 4a, with increased age, ERP individual mean peak amplitudes and latencies also increased. For the N200, increased amplitudes indicated attenuation of this peak with age, as they become less negative. For the P300, a positive going ERP, the increase with age indicates peak amplitude amplification.

**Figure 4.**
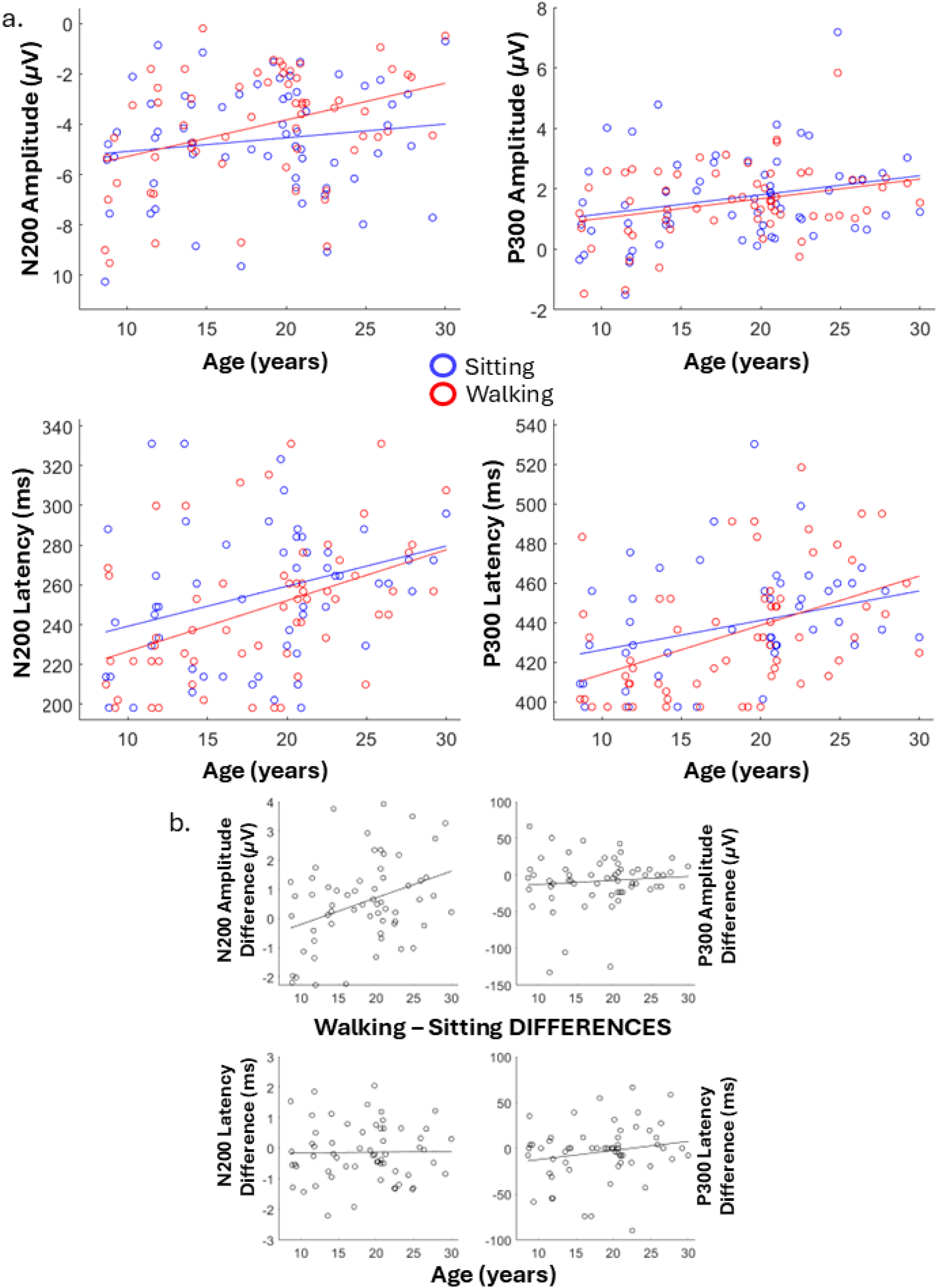
ERP peak amplitudes and latencies across age. A) On all four subplots, each point is a single participant’s mean peak N200 amplitude (upper left), N200 Latency (lower left), P300 amplitude (upper right), and P300 latency (lower right), plotted against their age. Blue markers are when sitting, and red markers are walking. Lines are the linear best fit for the data respective to that motion state. B) For all subplots here, the markers represent an individual participant’s difference between their walking mean and sitting mean for that plot’s respective measure, plotted against age. Lines are the linear best fit.

For the N200 peak amplitude, there was a significant interaction between motion state and age (*F*(1,59) = 8.864, *p* = .004, η_p_ ^2^ = .131). To further elucidate this interaction effect, we used linear regressions to function as *post-hoc* analyses. When sitting, the regression analysis indicated a non-significant relationship between age and N200 peak amplitude (R^2^ = .020, *F*(1,59) = 1.198, *p* = .278), but that was significant when walking (R^2^ = .140, *F*(1,59) = 9.630, *p* = .003). Figure 4b further visualizes this by showing the magnitude of the change in peak measure from sitting to walking for each individual as it relates to their age.

Grand average ERPs and scalp topography plots are presented in Figure 5a and 5b, respectively.

**Figure 5.**
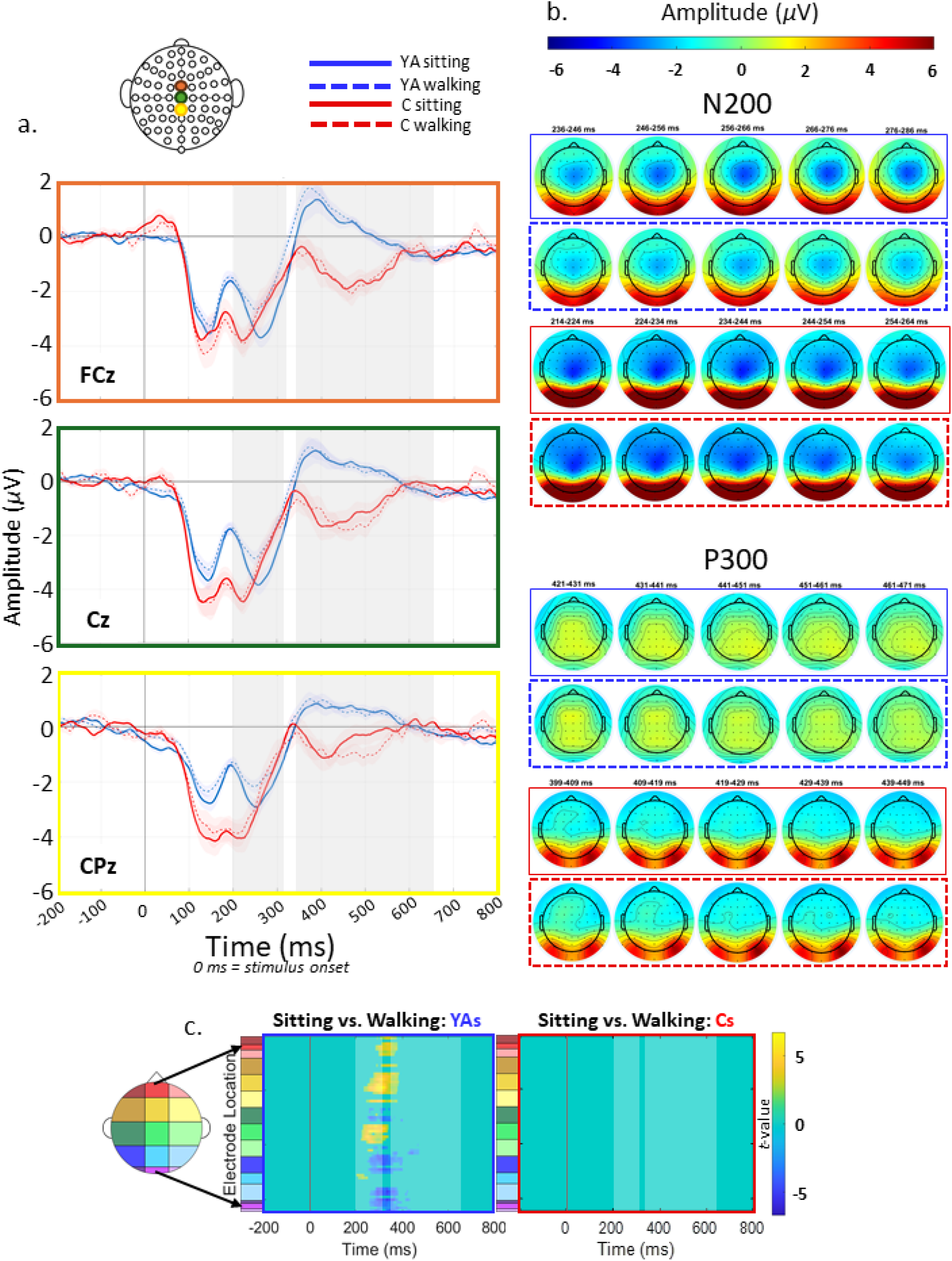
Neurophysiology by age group and motion state. A) Grand average ERP waveforms at FCz, Cz, and CPz. Border color of each waveform corresponds to the color of the marker on the scalp map at the top, to show electrode locations. 0 ms on the x-axis represents stimulus onset. Blue lines are Young Adults (YAs), red are Children (Cs). Solid lines are sitting, and dashed lines are walking, respective to their age group’s colors. Grey boxes highlight the latency window for the N200 (200-325 ms) and P300 (350-650 ms) ERPs. B) Grand average scalp topographies with latencies shown aligned to the peak of each age group’s respective average peak amplitude. Box borders correspond to the same color and type legend as in the ERP waveforms in A. C) Statistical cluster plots showing results from cluster-based permutation analyses examining the difference between sitting and walking within either age group.

Results of the cluster-based permutation analyses are visualized in Figure 5c. The young adults demonstrated clusters of statistical differences in neural activity between sitting and walking conditions. Most of these clusters occurred between approximately 200 – 450 ms after stimulus presentation. The effects that this approach revealed within this age group were ERP amplitude reductions, shown as yellow clusters, during walking over frontal and frontocentral scalp (red, and yellow on the scalp map on the far left), and continuing over central-parietal scalp (yellow and green). Blue clusters are seen at more parietal and occipital (blue and purple on the scalp map) locations, but also represent amplitude reductions when walking, though the t-value is negative because the amplitude values are positive. The children had no clusters of significant differences between sitting and walking.

### 3.4 Gait Kinematics

When separated by age group, there was a significant difference between chosen treadmill speed (*t* (59) = 9.860, *p* <.001), (Table 1). However, chosen treadmill speed significantly correlated with height (Pearson’s *r* = .730, *p* < .001). Treadmill speed differences have been shown in young adults not to significantly influence cognitive performance on a working memory task (Kline et al., 2014), nor ERP measures (De Sanctis et al., 2014). Therefore, height is used as a covariate in gait analyses to control for this and to assess developmental trajectory.

For the rmANCOVA on gait metrics, significant values are reported here along with partial-eta squared. If dual-task state, walking only or walking while performing the vGNG task, has a significant main effect, Bonferroni’s *post-hoc* pair-wise tests were performed. Results are visualized in Figure 6, and all effects are summarized in Table 2. Unsurprisingly, there was a significant effect of height for all gait measures: Stride Distance mean (*F*(1,55) = 97.567, *p* < .001, η_p_^2^ = 0.640) and variability (CV%) (*F*(1,55) = 57.150, *p* < .001, η_p_^2^ = 0.510); and Stride Duration mean (*F*(1,55) = 37.332, *p* < .001, η_p_ ^2^ = 0.404) and variability (CV%) (*F*(1,55) = 34.919, *p* < .001, η_p_ ^2^ = 0.388).

**Figure 6.**
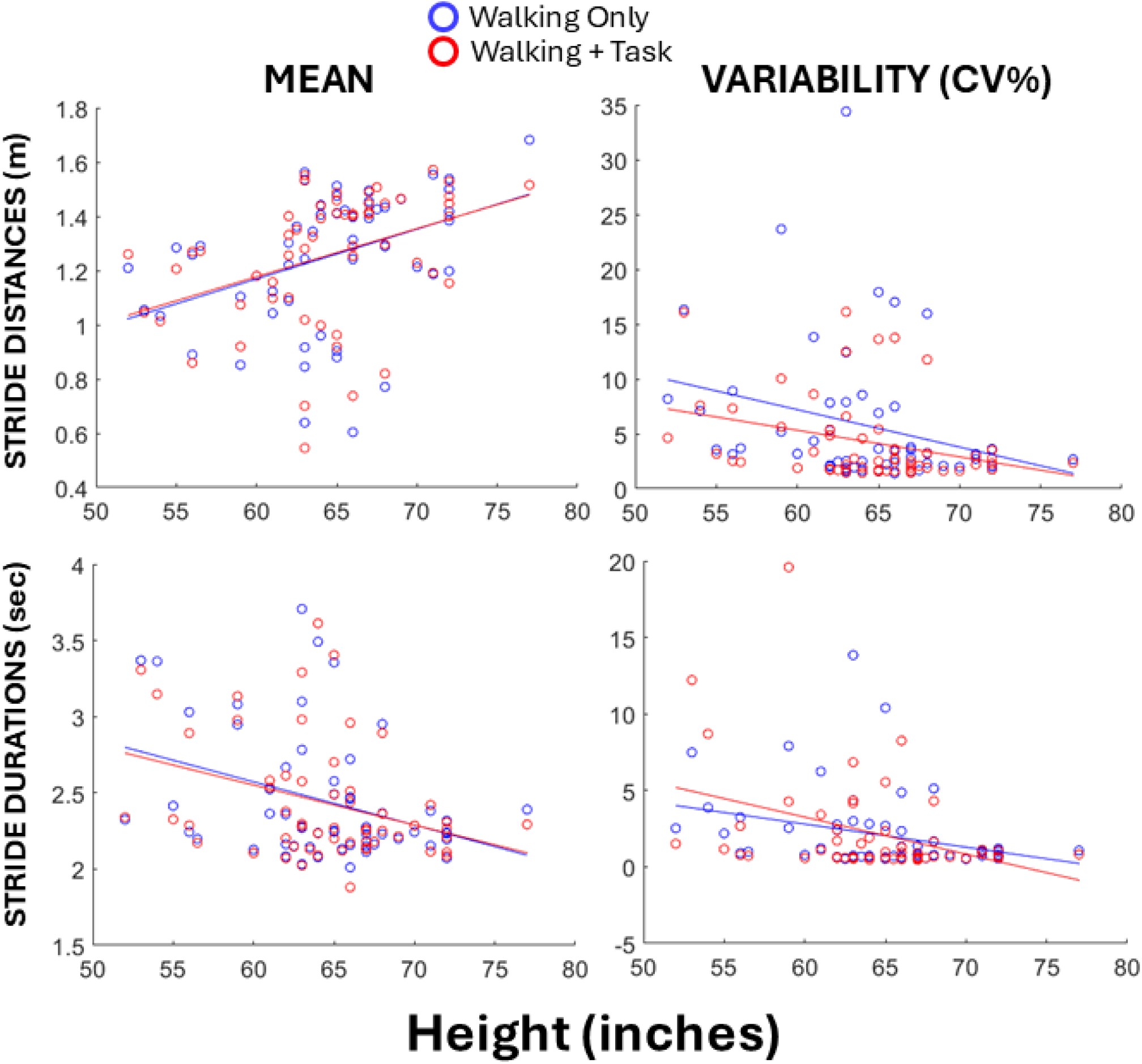
Single and dual-tasking gait kinematic measures in relation to participant height. All lines are linear best fit. On all plots, each marker represents a single participant. For each participant there is a blue marker for their measure when walking only, and a red marker for when walking while performing the vGNG task.

There was also a main effect of dual-task state on variability of stride distance (*F*(1,55) = 5.745, *p* = .020, η_p_ ^2^ = 0.095), and an interaction between dual-task state and height on variability of stride distance (*F*(1,55) = 4.702, *p* = .034, η _p_^2^ = 0.079). We used linear regression as *post-hoc* analyses to further characterize this interaction effect. The regression analysis showed both a significant relationship between height and variability of stride distance when walking only (R^2^ = .408, *F*(1,55) = 37.877, *p* < .001), and an even stronger relationship when dual-task walking (R^2^ = .567, *F*(1,55) = 72.164, *p* < .001).

## 4. Discussion

Using MoBI to investigate gait, cognitive performance, and neurophysiologic changes under multi-modal demands, we observed effects of cognitive-motor interactions that were related to age in a cohort of 61 neurotypical children and young adults. These may serve as developmental markers of multi-domain processing, integration, and execution. One of these was neurophysiologic: progressively larger attenuations of the N200 ERP peak amplitude between sitting and walking were observed with greater age. The second was motoric, with the largest reductions in stride distance variability from walking-only to dual-task walking being observed in participants with shorter heights. Table 2 and Figure 8 provide a summary of all findings.

We also observed several main effects of development, regardless of motion state. With increased age, N200 amplitude and latency, P300 amplitude and latency, vGNG task accuracy, and stride distance also increased. Reaction times also speeded up, with stride duration, and variability of stride distance and duration decreasing. One main effect of C-M interaction along the motor axis was observed: stride distance variability was greater when walking only, compared to walking and doing the cognitive task.

These results together provide an encompassing view of the changes to C-M interactions through childhood and into young adulthood and characterize a distinct pattern of effects that emerge during typical development.

### 4.1 Behavior

No main effect of motion state, sitting vs. walking, on either task accuracy or reaction time was observed. Increased participant age was significantly correlated to overall increased accuracy and faster reaction time. C-M interaction did not impact task accuracy or reaction time in this group of participants, indicating that – at least at the group level – neurotypical children and young adults are able to maintain cognitive task performance under C-M demands. Though, we do want to be careful to distinguish an absence of a significant behavioral effect from evidence of equivalent performance. The main influencing factor here was age, regardless of motion state. This is in line with established understanding of executive function development through childhood, adolescence, and into young adulthood. Although there was no main effect of motion state, individual participants showed increases or decreases in accuracy, reaction time, or both (Figure 2). This replicates results reported in previous research from our research group in young adults (De Sanctis et al., 2014; Patelaki et al., 2022), and extends the effect to those under 18 years of age. Future research could use a similar analysis approach as Patelaki et al., and separate EEG and gait analyses based on groups of those who did and did not improve accuracy when walking. The current results provide a critical foundational understanding against which to compare similar developmental trajectories in neurodevelopmental conditions (Lau-Zhu et al., 2019). A motion-state interaction effect with age, or a lack of any significant relationship with age within a particular clinical group may provide a marker for early-identification, sub-group phenotyping, or markers for identifying individuals who may particularly benefit from specific types of physical or occupational therapies, educational interventions, and other clinical decisions.

### 4.2 Neurophysiology

Age had a significant main effect on both N200 and P300 amplitudes and latencies, with all increasing with older age. Topographically, the loci of the ERPs also became more focused with age.

We observed that the N200 peak amplitude becomes less negative and attenuates with age. This reduction with age observed in the current study was expected based on previous literature, and many studies have reported latency reductions with age (Cragg et al., 2009; Davis et al., 2003; Downes et al., 2017; Hoyniak, 2017; Pires et al., 2014). However, the results here demonstrated a latency increase with age, which contrasts with previous literature (Enoki et al., 1993; van der Stelt et al., 1998). Though, the consistency of this is mixed in the literature, with others reporting little change in the N200 through childhood (Johnstone et al., 2007).

The P300 positive-going peak’s amplitude increases with age, demonstrating an amplification and this is consistent with the literature. However, the latency increase observed here is not (Downes et al., 2017; Fuchigami et al., 1995; Johnstone et al., 2007; Pires et al., 2014; Riggins & Scott, 2020; Rozhkov et al., 2009). It should be acknowledged, though, that the literature we are comparing the current results against are not from MoBI studies so it is reasonable that the developmental trajectories of C-M interaction-related ERPs might not exactly replicate, given the vastly different demands of a MoBI task. However, these effects were also observed in the more traditional sitting blocks, so C-M interactions cannot be the only explanation. Earlier peak latencies in ERP research are typically interpreted to reflect faster cognitive processing and greater neural efficiency. However, the earlier ERP latencies of our younger participants were accompanied by poorer task performance, which could reflect inefficient processing of the vGNG under single- and dual-task conditions. While not included in our original hypotheses and planned analyses, there is a visually observable difference between age groups in the amplitude of the P2 ERP component, which precedes the N200, with children having reduced amplitudes. The P2 component is related to attentional allocation and early stimulus encoding and has been suggested to influence subsequent ERP components like the N200 and P300, reflecting the extent of allocated attentional resources and stimulus information available for the subsequent higher order processes (Barry & De Blasio, 2013; Ghin et al., 2022; Stock et al., 2016). Essentially, younger brains may not have optimally engaged in the attentional conflict monitoring and earlier processing necessary for successful response choices, thus impacting the broader time-course of neural activity.

The finding that age modulates the effect of motion state (sitting versus walking) on N200 peak amplitude is novel. Increased age was significantly related to greater magnitudes of attenuation of the N200 amplitude when walking, but not when sitting. Our hypothesis was that greater C-M interference would be evidenced this way, by weaker ERP peak differences between sitting and walking in younger participants. This was accompanied by reduced accuracy and slower reaction times in younger participants, but that effect was not dependent on motion state so we cannot conclude that the greater neurophysiological adjustment in the older participants is tied to their overall better vGNG task performance and therefore reflective of strong cognitive-motor integration.

Younger participants made no observable adjustments in neural processing but maintained task performance when walking. Further, this was maintenance of slower and less accurate performance. It is possible that the younger participants demonstrated both worse performance with the vGNG task and an unrelated concurrent lack of available neural flexibility to adjust under dual-task demands. It is also possible that, 1) it was easy to maintain their lower and slower-performance, negating any need for reallocation of neural resources, whether they had access to the flexibility to do so or not, or, 2) there was a floor effect, so their performance couldn’t get much poorer when walking unless there was some fundamental misunderstanding of the task - though we are confident this wasn’t the case as study personnel confirmed understanding throughout the experiment, and all participants performed above chance (d’ = 0) when both sitting and walking. On the other end of the age range, however, the older participants made significant neural adjustments in addition to the maintenance of their performance, which was overall faster and more accurate. This is a more challenging result to achieve and therefore their performance maintenance likely necessitated an adjustment and redeployment of the involved cognitive processes when walking, which is thus reflected in their significantly reconfigured neural activity patterns when compared to sitting.

The presentation of activity during the typically understood P300 latency window in the children is unusual. A P300 typically presents at central-parietal scalp locations as a positive-going and often prolonged peak from 300-600 ms after stimulus presentation and about 100 ms after the N200 peak. In the children we observed here, the activity does begin similarly within that latency window, beginning with a positive waveform about 100 ms after the N200 peak that peaks at about 350 ms, but it then transforms into another negative going, slow waveform. We see this negative deflection at all three central electrodes plotted in Figure 5 (FCz, Cz, and CPz), reaching maximal negativity at around 450-500 ms before the waveform potential becomes positive-going once more, and gradually returns toward baseline through the remainder of the epoch. The scalp topography plots during this time window as shown in Figure 5 are also remarkably different between the age groups. This atypical presentation of the P300 in the children makes it difficult to interpret the results of our *a priori* analytical approaches as they were contextualized under the expectation of a more typical NoGo P300 waveform. Occasionally, the P300 is divided into the P3a and P3b sub-components, even in children (Fauvel, 2025; Peisch et al., 2022). However, this doesn’t seem to apply to the data here, because the children do not demonstrate a second positive peak after this negativity that would be demonstrative of a P3a-P3b component, nor is this subdivision observed in other adult MoBI vGNG studies (De Sanctis et al., 2012; De Sanctis et al., 2014; De Sanctis et al., 2020; Malcolm et al., 2019; Patelaki, Foxe, Mantel, et al., 2023; Patelaki et al., 2022).

This waveform could also be compared to the N450 ERP. The N450 is a slow negative waveform that is mid-frontally located, onsets around 400-600 ms after target stimulus presentation. It is often observed in Stroop-like tasks as a marker of conflict, and is thought to be generated by the anterior cingulate cortex (Gibbons, 2011; Hseih et al., 2018; Liu et al., 2014; Szűcs & Soltész, 2012). In one study it has been observed in neurotypical children, with kids having increased amplitudes compared to young adults (Larson et al., 2012). To our knowledge, though, the N450 has not been reported with Go/NoGo tasks and therefore it may be too speculative to interpret our observations as reflecting a novel version of the N450.

Another possibility for interpretation is the classical N400 component (Toffolo et al., 2022, 2024). This ERP is triggered by violations of expectation, particularly ones related to semantic processing. Stimuli in N400 paradigms have semantic features and can be visual or auditory but are not necessarily always words or sentences (Hamm et al., 2002; Kutas & Federmeier, 2011). Previous studies have used semantically congruent (e.g., spoon and fork) and incongruent (e.g., spoon and tree) photos to elicit an N400, with the incongruent pairs yielding a larger N400 amplitude broadly distributed over central scalp (Dini et al., 2022; Hamm et al., 2002; Petrusic et al., 2021; Toffolo et al., 2022, 2024). It is possible that the children in this study deployed some form of semantic processing as a strategy to the working memory facet of performing the vGNG task used herein. As participants were older, this strategy was less necessary to accomplish the demands of the task, manifesting as the progressive loss of this unexpected negative ERP we observed.

This ERP could serve as another marker of C-M interactions in typical development that is independent of motion state, though more research would be needed to validate this observation. The observation of this component appeared independent of motion state, though this is only based on exploratory visual inspection. Its presence in the children may contribute to why a motion state effect for the P300 was not observed, due to the parameters chosen *a priori* for these analyses such as the selected time window and voltage deflection. However, the children still had no clusters of significant difference in the *post-hoc* exploratory cluster analyses. These data are a valuable addition to the interaction effects of N200 amplitude and stride distance variability that we observed. Having an understanding of an additional marker that is specifically elicited under these recording methods and is unaffected by motion state can provide another way to identify groups who differ in C-M interactions across development. For example, future research may hypothesize that a particular clinical group, under dual-task demands, would demonstrate an atypical effect of motion state on this component’s peak amplitude, a weaker relationship with age, or both.

We visualized the peak amplitude of this explorartory component and its relationship to age in Figure 7 for reference and future hypothesis generation. Similar to our approach to the N200 and P300 peaks, we calculated the minimum value (most negative amplitude) in the 400-600 ms window of the epoch at FCz for each participant, based on visual inspection of the grand average ERPs of the children. We strongly emphasize that the presentation of these data should be considered exploratory only, and that this novel component and our conservative interpretation of it require further investigation.

**Figure 7.**
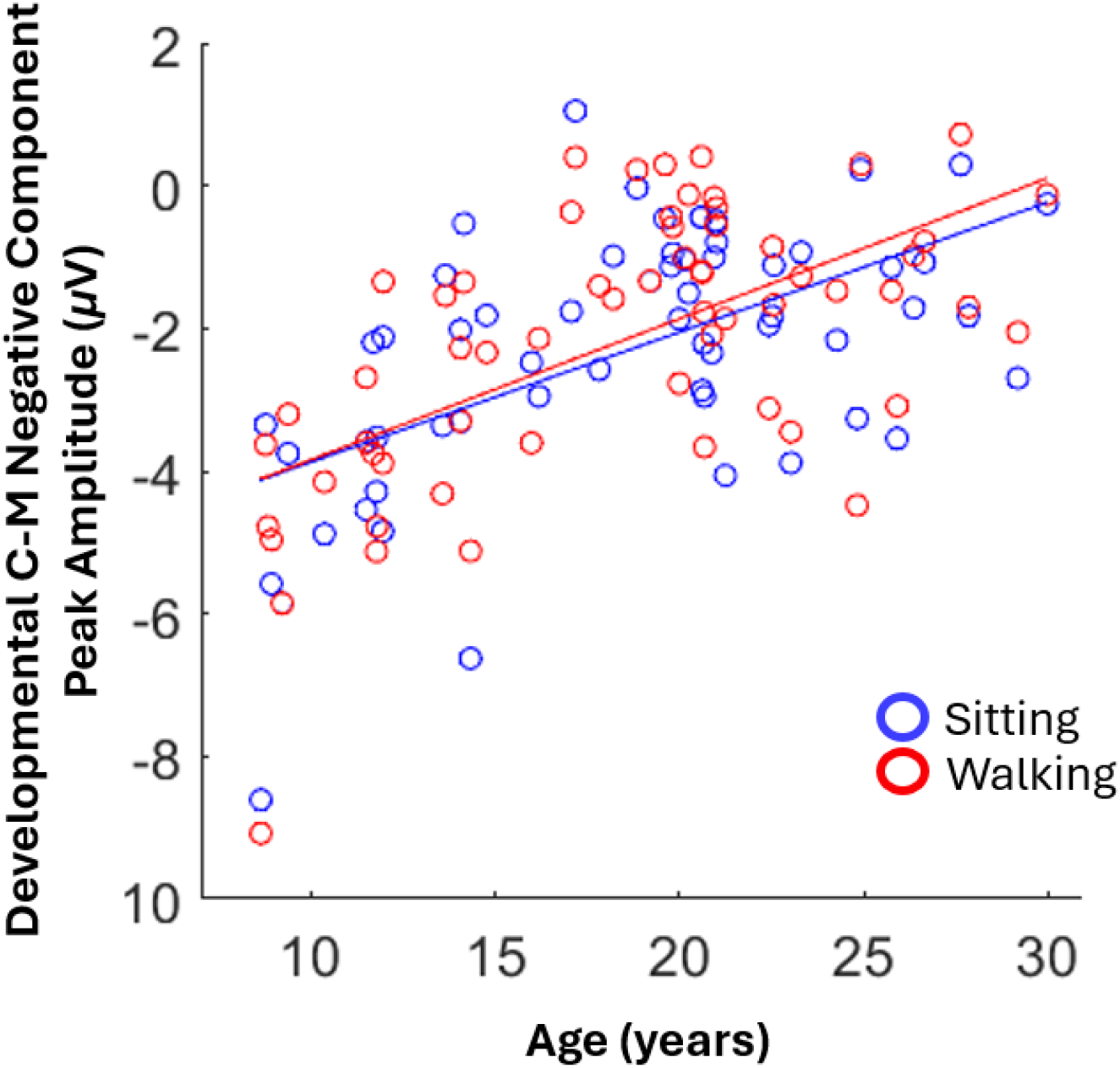
Scatter plot of participant age and corresponding mean peak amplitude at FCz of the exploratory developmental C-M negative component observed in the data. Each marker is a single participant’s mean respective to each of the two motion states: blue for sitting and red for walking.

**Figure 8.**
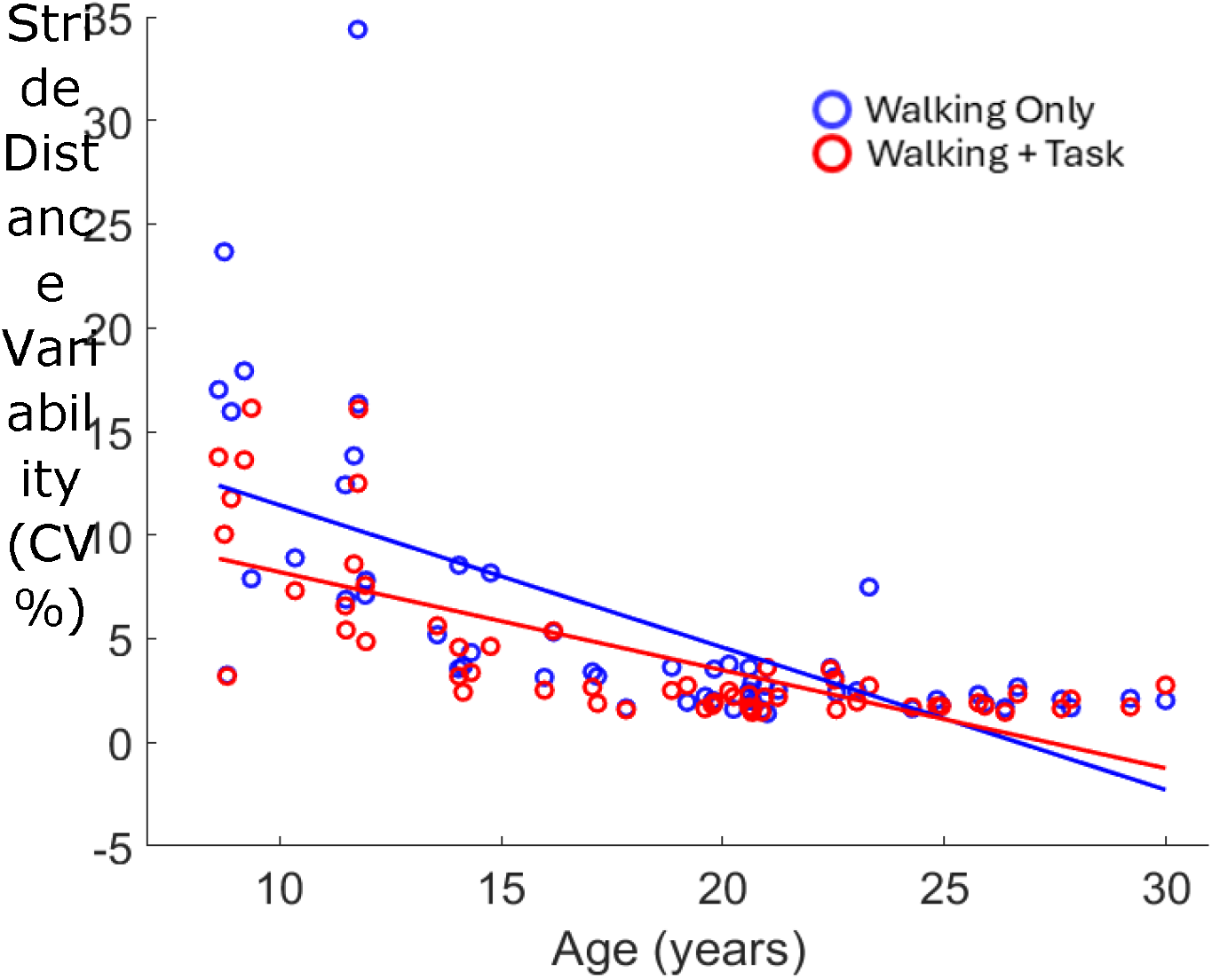
Scatter plot of participant age and stride variability (CV%). Each marker is a single participant’s mean respective to each of the two motion states: blue for walking only and red for walking and doing the vGNG task.

### 4.3 Gait Kinematics

The data demonstrated a main effect of cognitive task state on stride distance variability, which replicated what has previously been observed in neurotypical young adults under similar MoBI study designs (Patelaki et al., 2022; Richardson et al., 2022) and then further extends this knowledge into younger ages. Previous interpretations of this effect posit that since participants were walking on a flat non-obstructed surface at a constant speed, an automatic (hence, less variable) gait pattern might be more easily adopted, thus freeing cognitive resources to address other task demands.

The interaction between cognitive task state with height is novel and demonstrates that the dual-task effect is strongest at younger ages and shortest heights, and this observation is supported by previous research (Kraan et al., 2017; Oudenhoven et al., 2019). Prior work has shown that stride-to-stride variations and variability decrease through ages as young as 3 to 14 years (Hausdorff et al., 1999). Increased variability in younger ages has been suggested to be due to reduced postural stability at slower treadmill walking speeds (Zijlstra et al., 1996), immature physical and neural control systems (Fairley et al., 2010), and less efficient integration of movement with visual and proprioceptive information (Holst-Wolf et al., 2016; Mallau et al., 2010; Müller et al., 2013). These characteristics may explain why children showed greater stride distance variability during walking alone, leaving more room for variability to decrease when they also performed the cognitive task. Older/taller participants may already walk with a more constrained and stable gait pattern, leaving less opportunity for variability attenuation when shifting from walking only to dual-task walking.

Further, height was a better predictor of stride distance variability when walking and doing the vGNG task, compared to walking only, indicating that the C-M interaction here may provide a valuable marker of the development of C-M integration that may be better than gait measures when walking alone. This may provide a more sensitive measure derived under more naturalistic experimental settings via MoBI that can be applied to future research in typical and clinical groups.

Of note, when visually inspecting the relationship between stride variability and height (Figure 6, upper right), it was observed that there was a cluster of individuals with higher (approximately 5-20 CV%) variability but mid-range (approximately 60-70 inches) heights. While we chose height as the covariate in this analysis based on the presumption that it would most appropriately control for differences in participants physical bodies and individual stride lengths, this choice may not have captured a potential effect of gross motor development – specifically, children who are taller but younger. These kids may have more variable locomotion due to underlying motoric developmental progress, not overall height. To visualize this exploratory hypothesis, we plot in Figure 8 the relationship between age and stride variability in the same manner as was done with height. Preliminarily, this seems to more robustly depict the change in stride variability through development.

### 4.4 Limitations

It’s important to acknowledge that the C-M interactions we observe here may present differently under different experimental design choices such as task difficulty (Patelaki, Foxe, McFerren, et al., 2023), task executive function or domain (Lim et al., 2025; St George et al., 2022), motor demand type or complexity (Downey et al., 2022; Liu et al., 2024), and others. More MoBI studies are needed to confirm if the results we observed are reliable. From a development perspective, the cross-sectional design of the current study provides a good foundational understanding but results from future longitudinal studies would be more powerful. Additionally, this study formulated hypotheses and methods focused on successful exhibitions of C-M interactions, correct-rejections. It is possible this set of results can be further contextualized using error-related approaches.

Another possible limitation is the use of a treadmill and participant-selected treadmill speed. The predictability inherent to the treadmill is beneficial to experimental design and analysis, but fixed-speed walking is less demanding than over-ground or self-paced treadmill walking (Kao & Pierro, 2021). We acknowledge that using a treadmill adds an element of external motivation to walking control, as opposed to over-ground walking which is internally motivated and controlled in everyday life. However, the treadmill’s ability to maintain a consistent motor load (speed) throughout the entire experiment is valuable in complex study designs such as with MoBI (De Sanctis et al., 2014; Malcolm et al., 2018; Richardson et al., 2022).

Some of the younger children had limited experience with a treadmill. Observationally, the study team believes that some of the children may have been underestimating their walking ability when choosing a treadmill speed, but once it was chosen, we did not want to change it because all walking blocks were to be performed at the same walking speed. This may have contributed to the increased variability of gait in the children for a couple of reasons: 1) walking slowly has been shown to be, at least in adults, a form of motor demand compared to a natural or “usual” pace (Almarwani et al., 2016). And, 2) as the children walked some would occasionally ‘play’ on the treadmill. Examples of this included walking to the front-most point of the treadmill, standing still, and then letting the belt move them backwards to the center, at which point they would start walking again; walking in “silly” ways such as taking marching or skipping-like steps; or walking in a laterally wavy pattern instead of straight ahead. As admittedly entertaining as these behaviors by the children were, study personnel were attentive to these and would give verbal corrections (i.e., “Please walk as normally as possible.”) whenever they were observed in a participant. Further, as mentioned in the methods, our processing pipeline excluded gait cycles with trajectories with a kurtosis greater than 5 standard deviations of the mean, so cycles during these types of behaviors were likely rejected from analysis. To address these limitations, future research could explore ways to pre-determine ‘optimal’ walking speed that takes into account individual factors like height and age, to remove some of the subjectivity from the decision.

## 5. Conclusion

Our main goal was to investigate how C-M interactions present across development in a neurotypical population using MoBI. Several main effects of development (age or height) were found, as shown in Table 2, but two interaction effects emerged that may serve as novel developmental markers of C-M interactions. The magnitude of attenuation of the N200 amplitude under dual-task conditions increased with age, and dual-task attenuation of stride length variability decreased with age (Figure 9). A third possible marker presents as a novel later ERP waveform present only in younger ages, which we’ve called the dCM-Nc, that appears to be specific to our methods but unimpacted by motion state. These markers have potential for use in research to further elucidate factors that influence their trajectories in neurotypical development. Practice effects (Strobach, 2020) or individual differences may influence C-M interactions to be more interfering or integrative for a given group. Further, this foundational knowledge can be applied to research in neurodevelopmentally divergent populations to learn if and how these markers differ and utilize them to possibly identify phenotypic subtypes within clinical groups, promote early or more accurate detection and diagnosis, or even differentiate individuals who may particularly benefit from specific therapeutic or educational interventions.

**Figure 9.**
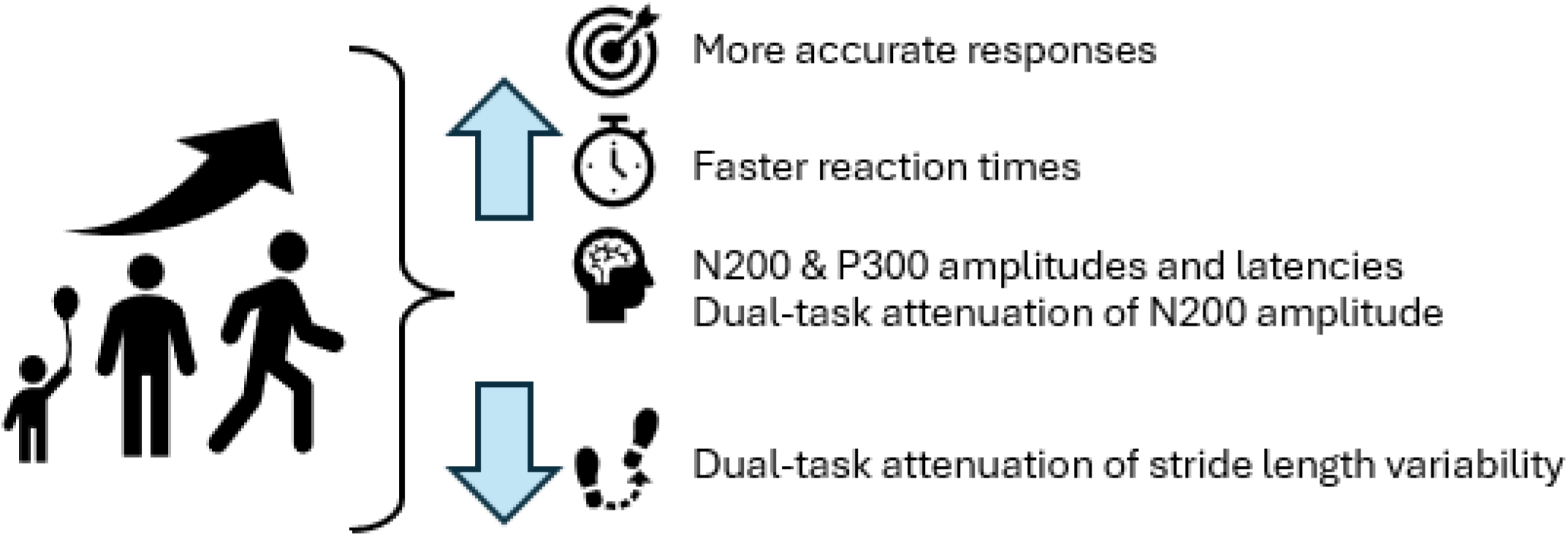
Visual Summary of Results

## DECLARATIONS

### Ethics and Consent

The Research Subjects Review Board of the University of Rochester approved all the experimental procedures (STUDY00006574). Each participant provided written informed consent in accordance with the tenets laid out in the Declaration of Helsinki.

## Acknowledgements

We sincerely thank our participants and their family members for taking part in this research. Special thanks to Dr. Eleni Patelaki, Ph.D., Dr. Kathryn Toffolo, Ph.D., and Ms. Eve Lang. The authors also acknowledge the support of the entire research team at the Frederick J. and Marion A. Schindler Cognitive Neurophysiology Lab.

## CRediT Author Contribution Statement

Paige R. Nicklas: Conceptualization, Data curation, Formal analysis, Investigation, Visualization, Writing – original draft, Writing – reviewing & editing. **Edward G. Freedman**: Conceptualization, Formal analysis, Investigation, Supervision, Resources, Visualization, Writing – original draft, Writing – reviewing & editing **John J. Foxe**: Conceptualization, Formal analysis, Investigation, Supervision, Resources, Visualization, Writing – original draft, Writing – reviewing & editing

## Data Sharing

The stimuli, supporting datasets, and code for this experiment can be made available from the corresponding authors on reasonable request.

## Consent for publication

Not applicable.

## List of Abbreviations

AMICA: Adaptive mixture independent component analysis
ANCOVA: Analysis of covariance
C-M: Cognitive-motor
CPz: Midline centroparietal electrode
CR: Correct rejection
Cs: Children
CV: Coefficient of variation
CV%: Coefficient of variation expressed as a percentage
Cz: Midline central electrode
dCM-Nc: Developmental cognitive-motor negative component
DT: Dual-task / dual-tasking
EEG: Electroencephalography
ERP: / ERPs Event-related potential / potentials
FCz: Midline frontocentral electrode
FSIQ: Full-scale intelligence quotient
IAPS: International Affective Picture System
IC /: ICs Independent component / components
ICA: Independent component analysis
IQ: Intelligence quotient
LSL: Lab Streaming Layer
MoBI: Mobile brain-body imaging
rmANCOVA: Repeated-measures analysis of covariance
RT: Reaction time
SD / STD: Standard deviation
SPSS: Statistical Package for the Social Sciences
ST: Single-task / single-tasking
vGNG: Visual Go-NoGo
WASI-II: Wechsler Abbreviated Scale of Intelligence, Second Edition
WO: Walking-only
YAs: Young adults

